# Improving Timeliness in the Neglected Tropical Diseases Preventive Chemotherapy Donation Supply Chain through Information Sharing: A Retrospective Empirical Analysis

**DOI:** 10.1101/2021.06.03.446886

**Authors:** Elena Kasparis, Yufei Huang, William Lin, Christos Vasilakis

**Affiliations:** Department for Health, University of Bath, Bath, United Kingdom; Trinity Business School, Trinity College Dublin, Dublin, Ireland; School of Management, University of Bath, Bath, United Kingdom

## Abstract

**Background:** Billions of doses of medicines are donated for mass drug administrations in support of the World Health Organization’s “Roadmap to Implementation,” which aims to control, eliminate, and eradicate Neglected Tropical Diseases (NTDs). The supply chain to deliver these medicines is complex, with fragmented data systems and limited visibility on performance. This study empirically evaluates the impact of an online supply chain performance measurement system, “NTDeliver,” providing understanding of the value of information sharing towards the success of global health programs.

**Methods:** Retrospective secondary data was extracted from NTDeliver, which included 1,484 shipments for four critical medicines ordered by over 100 countries between February 28, 2006 and December 31, 2018. We applied statistical regression models to analyze the impact on key performance metrics, comparing data before and after the system was implemented.

**Findings:** The results suggest information sharing has a positive impact on three performance indicators: purchase order timeliness (β=1.01, p<0.000), arrival timeliness (β=0.53, p=0.09), and—most importantly—delivery timeliness (β=0.73, p=0.03). Three variables indicated an increased positive impact when the data is publicly shared: shipment timeliness (β=2.57, p=0.001), arrival timeliness (β=2.88, p=0·003), and delivery timeliness (β=2.82, p=0.01).

**Conclusions:** Our findings suggest that information sharing between the NTD program partners can help drive improved performance in the supply chain, and even more so when data is shared publicly. Given the large volume of medicine and the significant number of people requiring these medicines, information sharing has the potential to provide improvements to global health programs affecting the health of tens to hundreds of millions of people

**Author Summary:** The supply chain to deliver donated preventive chemotherapy medicines is complex due to the many stakeholders and partnerships participating, as well as challenging because the logistics are further complicated by delivery to remote destinations in developing countries. As MDA campaigns involve treating hundreds of thousands to millions of patients in endemic regions within entire countries over the course of days or weeks, close coordination and timing of medicine delivery is critical. Inefficiencies caused by fragmented data systems and limited transparency on supply chain performance further challenges the ability to identify shipment issues and explore the root cause of the issues. Prior to 2016, delivery was performing below standards, lagging as much as 40% below the WHO target of 80% on-time delivery. These delays result in wasted medicine donations, increased program costs, delayed MDAs, or sometimes even completely missed MDAs. In September 2016, an online supply chain performance measurement system, “NTDeliver,” was launched by the NTD Supply Chain Forum (a public-private partnership focused on managing and improving the PC donation supply chain) to enhance supply chain performance and information transparency. Our findings suggest that information sharing through NTDeliver is positively associated with performance at several key stages in NTD supply chain and that information sharing has more substantial positive impact on performance when the information is made publicly accessible, focused towards country program managers. The study findings support investment in supply chain systems and commitment to data transparency, in the context of a growing focus on supply chain investment in NTD programs.

## Introduction

Public-private partnership programs provide medicines for preventive chemotherapy (PC) through mass drug administration (MDA) campaigns to more than one billion people annually. The programs are sustained by large-scale donations from major pharmaceutical companies in support of the World Health Organization’s 2012 “Roadmap to Implementation,” which outlined global strategies and 2020 targets to control, eliminate, and eradicate Neglected Tropical Diseases (NTDs) [1]. While significant progress was made towards these 2020 targets, the WHO has recently released a new NTD roadmap with 2030 targets, which pharmaceutical manufacturers have committed to continuing to support [2]. As of January 2020, 15 billion doses of medicines were donated towards these PC-NTD programs [3]. The donations from pharmaceutical companies are what makes these the world’s largest and most successful public health programs [4]. MDA campaigns are comprised of once or twice-a-year treatment with one or more medicines at the community level that bring together a number of stakeholders, requiring considerable coordination as they typically involve treating hundreds of thousands to millions of patients in endemic regions within entire countries over the course of days or weeks [5].

The logistics involved to make these medicines available to support MDAs is both critical and complex, due to the importance of meeting the targeted treatment date and the many stakeholders and partnerships involved. Fig 1 provides an overview of the processes involved and performance measures associated with each link in the supply chain. For the MDA campaigns, there are considerable resources and coordination involved within a narrow timeframe, which thereby increases the criticality of on-time delivery. However, inefficiencies caused by fragmented data systems and a lack of visibility to supply chain performance have resulted in substandard performance for on-time delivery of medicine to in-country central medical stores. Sometimes delivery targets may lag as much as 40% below the WHO target for 80% of all shipments to be delivered at least one month before the planned MDA date [6]. Delivery delays result in waste, increased program costs, and delayed or even completely missed MDAs, leaving individuals susceptible to NTD infections [7].

**Fig 1.**
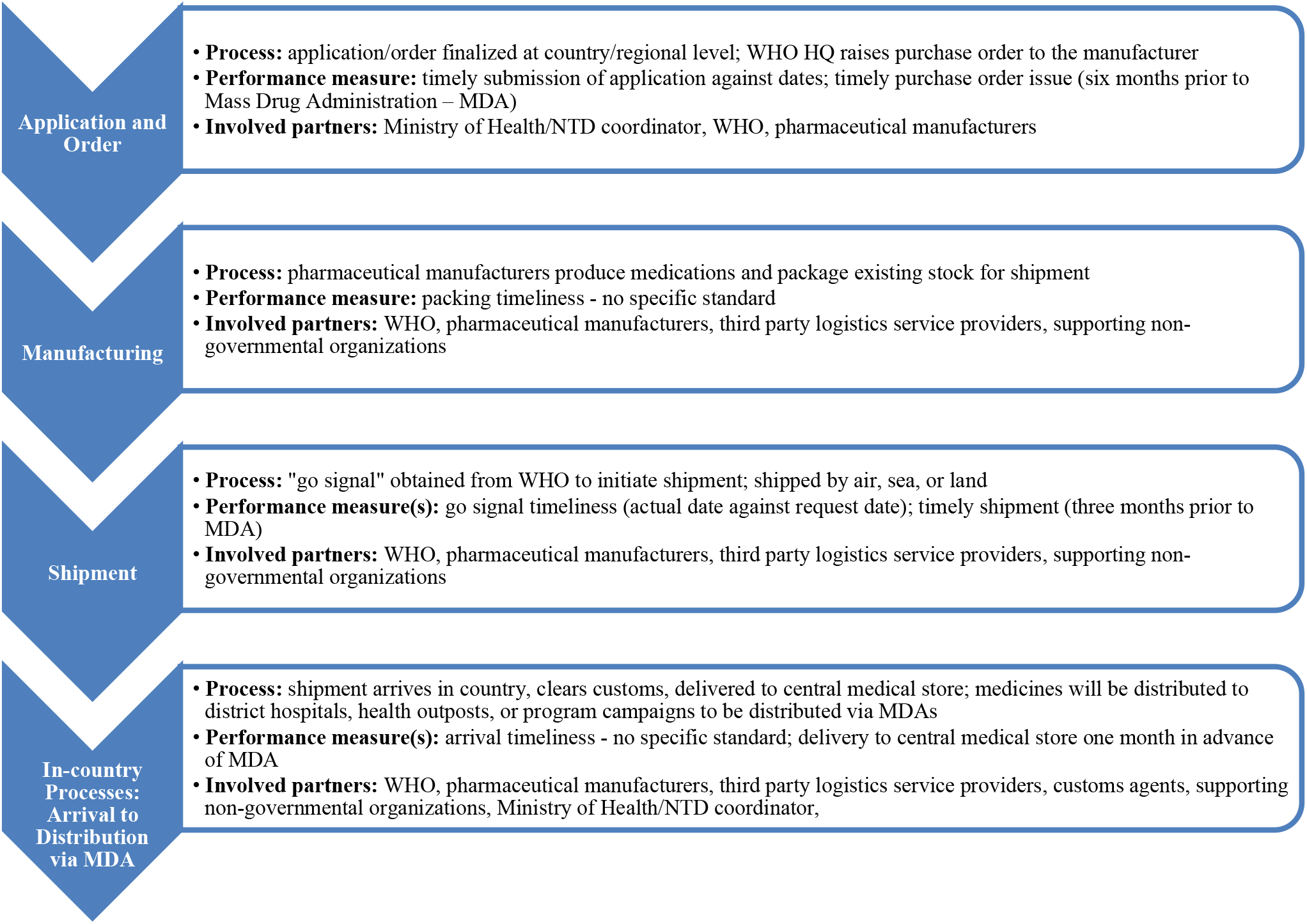
Components of the NTD supply chain from application for donation to in-country delivery to central medical stores.

To improve the efficiency and performance of NTD supply chain, the NTD Supply Chain Forum was established in 2012 [3]. The NTD Supply Chain Forum includes the following partners: the WHO, the Bill and Melinda Gates Foundation, six pharmaceutical donors (GlaxoSmithKline, Merck & Co. Inc., Merck KGaA, Pfizer, Johnson & Johnson, and Eisai), logistics partner DHL, and non-governmental organizations (NGOs) - the International Trachoma Initiative, Children Without Worms, the Mectizan® Donation Program, and RTI International [3]. Subsequently to the formation of this forum, “NTDeliver,” a centralized information system, was launched in 2016 by the NTD Supply Chain Forum to share data from various partners along supply process chain as means of facilitating performance information sharing [8]. Through NTDeliver, all stakeholders in the supply chain—and even the general public—may access performance metrics on all shipments of the four medicines that participate in data sharing through NTDeliver.

Information sharing in the context of supply chains— “the extent to which crucial and/or proprietary information is available to members of the supply chain”—is an integral aspect of performance management and the sharing of accurate and timely information has been linked to supply chain performance improvements [9,10]. Many studies have been conducted on the value of information sharing to improve supply chain performance in the private sector. Information sharing has been proven to have a range of benefits, from improved resource utilization to reduced cycle time between order and delivery [11]. These benefits stem from the increased transparency that enable risks to be anticipated and shared among supply chain partners, which strengthens coordination to achieve optimal operational performance [11–13]. Largely, this body of literature with empirical studies investigates the value of information sharing in a commercial supply chain focused on a dyadic relationship, primarily between two partners, a buyer and a supplier [14].

Despite the growing importance of supply chain initiatives in the global NTD agenda, there is limited research dedicated to exploring measures to improve NTD supply chain performance nor the impact of information sharing. This limitation may be an indicator that performance measurement and management systems have not been widely developed and systematically implemented as part of the overall humanitarian supply chain strategy [15]. Only Korpoc’s 2015 research on the impact of the NTD “first mile” processes (the segment of the NTD supply chain covering up to delivery to central medical stores) on MDA timeliness acknowledges this area of NTD supply chain performance measurement by identifying the need for performance indicators and outlining suggested metrics [7]. Furthermore, the recent COVID-19 pandemic has raised the profile both of the criticality of publicly sharing timely data in the global health domai^n^ and the topic of assuring robust supply chains to meet global health goals [16–21]. Thus, the timing could not be better to study information sharing and supply chain management in the wider global humanitarian health context.

This study seeks to evaluate empirically the impact of information sharing via NTDeliver on supply chain performance—improvements which ultimately contribute to achieving the global NTD targets. We examine the following two research questions: 1) what is the impact of information sharing through NTDeliver on the performance of the NTD PC medicines donation supply chain? and 2) what is the impact on the performance when country-level data is made publicly accessible? We use data obtained from the NTD Supply Chain Forum and implement regression models on these research questions. We find that information sharing has a positive impact on three performance indicators of the NTD supply chain: purchase order timeliness, arrival timeliness, and delivery timeliness. Furthermore, when country-level information sharing is made publicly accessible, more substantial impact is observed primarily on the downstream indicators, with positive impact observed on shipment timeliness, arrival timeliness, and delivery timeliness.

## Methods

### Study design and scope

We used retrospective data from NTDeliver that are routinely collected from and managed by supply chain partners supporting delivery of PC medicines to central medical stores. Permission was granted to use this data by the NTD Supply Chain Forum. The data is derived from vetted, existing data sources managed by NTD supply chain partners, such as:

- WHO Preventive Chemotherapy and Transmission databank
- Data provided to WHO country offices by the countries’ Ministry of Health through the joint application process to request donations
- Purchase orders raised by the WHO headquarters
- Shipping documents generated by logistics service providers, in partnership with pharmaceutical donors

The data represents shipments of four medicines from four manufacturers to treat three different diseases, accounting for almost 11.5 billion doses of PC medicines to 103 recipient countries covering 1,484 total shipments from February 28, 2006, to December 31, 2018. The data is refreshed and uploaded from these various sources daily [22]. While there are numerous medicines for NTDs donated by various pharmaceutical manufacturers for the NTDs, this research’s scope focuses on PC medicines donations managed by the WHO through the “joint application package” (JAP) established in 2013, which supports an integrated review and subsequent reporting on medicines usage.^23^ The JAP streamlines the application for donation of multiple medicines, especially as medicines are co-administered where diseases are co-endemic [23]. These PC medicines include: diethylcarbamazine citrate, albendazole, mebendazole, and praziquantel [24]. This focus is justified by the considerable volume of medicines, the unique nature of this supply chain that includes WHO involvement, the importance of PC to achieve NTD targets, and accessibility of data through NTDeliver. There are opportunities to improve processes across the supply chain but the focus of this research will be on the segment from application to delivery to central medical stores, also referred to by partners as the “first mile” [7]. We chose to focus on the first mile due to the WHO drive to improve on-time delivery to central medical stores, the accessibility of relevant data, and opportunity to leverage information sharing among the many partners involved in this segment. Central medical stores (CMS) (most commonly utilized in Africa, Asia, and Latin America) serve as a warehouse and administrative facility that receives, stores, and manages medical supplies for national health programs and initiatives and are generally leveraged for humanitarian stock.^6,25^ Improving the on-time delivery of PC medicine to the CMS is critical to the downstream in-country distribution to program sites where they are needed [6].

### Variables

The variables for this analysis are various key performance indicators (KPIs), which are actively reviewed through the NTD Supply Chain Forum and of interest to the supply chain partners, including the WHO. The most critical KPI is delivery timeliness, by which the WHO evaluates performance of this first mile of the NTD supply chain [6]. Both the independent and dependent variables were created from the data in the system, reflecting the KPIs and benchmarks standards tracked by the NTD Supply Chain Forum partners. Table 1 summarizes the key variables in the analysis.

**Table 1.**
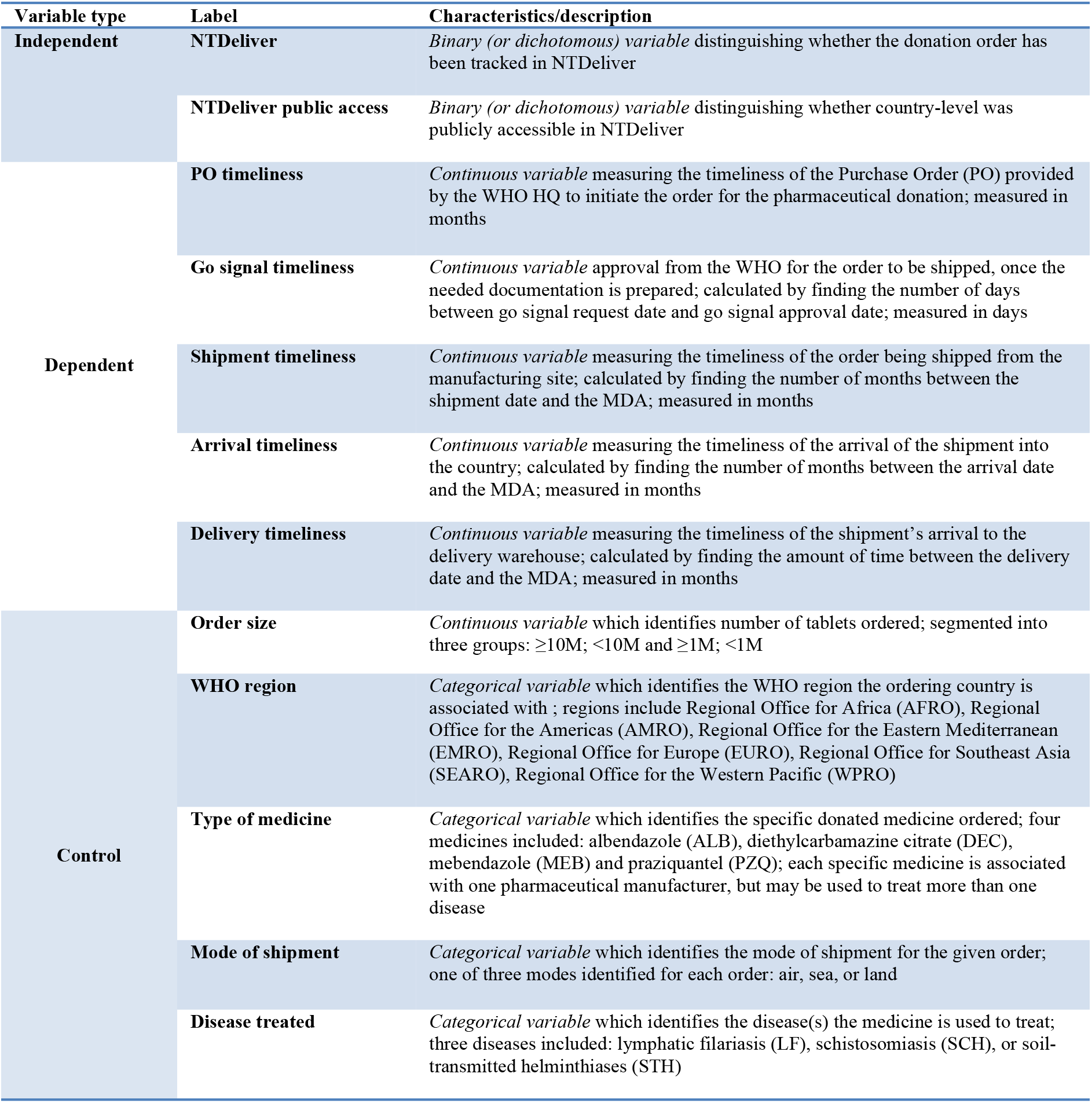
Variable definitions.

Confounding factors, incorporated via control variables, were included in the analyses to explore whether the relationship of the independent and dependent variables is skewed or invalidated by other confounding factors. The controls for medicine and disease were chosen with consideration to the fact that the medicines are produced by different manufacturers for different disease programs, which may lend itself to some variability in the supply chains. The region was also included as a control since performance may vary according to the destination – shipment routes and customs clearance processes vary according to the destination. Mode of shipment was incorporated as a control since the shipment speed varies between air, land, and sea. Finally, although the supply chain process is the same regardless of order size, larger orders could take more time to prepare and smaller orders are often shipped by air, making this also a necessary factor to control.

### Statistical analysis

A quasi-experiment design, using a “one-group pretest-posttest design without control group,” was chosen as NTDeliver was implemented in a real-world application that did not roll out the system in a phased approach, barring any ability to conduct randomization. This design was used to leverage historical data available on performance to understand the impact of this intervention. An ordinary least squares (OLS) regression model was used to review the relationship between implementing NTDeliver and its impact on delivery timeliness and other KPIs. Furthermore, the data meets the normality of the error distribution assumption for OLS regression. Q-Q plots were used to verify that most data is on a distribution lying on approximately on a straight line. While go signal timeliness did not show a straight trend, the central limit theorem enables the normality assumption to be met in the case of a “sufficiently large sample,” for which the literature generally notes a sample of >50 would be “robust to violation of the normality assumption” [26]. This variable had over 200 samples, thus meeting the central limit theorem condition. While both medicine and disease type are included as control variables, these variables are not found to be significantly correlated since some medicines treat more than one disease and therefore is not a 1:1 mapping; hence, multi collinearity is not a concern. Control variables and robustness checks were applied to strengthen the validity of the results. We considered p values less than 0.10 to be statistically significant.

The shipment data extracted from the system covers orders made through December 31, 2018 and was therefore segmented in two groups to address the first research question: 1) shipments with POs raised prior to the implementation of NTDeliver (February 28, 2006-August 31, 2016) for the “pre” NTDeliver group; 2) shipments with POs raised after the implementation of NTDeliver (September 1, 2016-Dec 31, 2018) for the “post” NTDeliver group. Only the data in the “post” group was used to answer the second research question regarding the impact on shipment performance of making country-level data publicly accessible. The data in the “post” group was split into two groups, with consideration to February 1, 2018, as the implementation date of this publicly accessible data: 1) “Post 1” = 0 for shipments with a PO date prior to February 1, 2018 but after August 31, 2016; 2) “Post 2” = 1 for shipments with a PO date equal to or later than February 1, 2018 but earlier than January 1, 2019.

## Results

We first study the general impact of information sharing through NTDeliver on shipment performance within the NTD PC medicines supply chain. The data collected included 1,484 total shipments, with 1,068 shipments classified in the “pre” NTDeliver group and 416 in the “post” group. As noted, pairwise deletion was used in cases where data was missing, which accounts for the differing number of observations between variables. Table 2 summarizes the regression results illustrating the bivariate associations between the implementation of the NTDeliver system and NTD supply chain KPIs.

**Table 2.** **Regression results showing the relationship between NTD supply chain key performance indicators and implementation of information sharing through NTDeliver**.

| Dependent Variable | PO timeliness<br>(in months) | Go signal timeliness<br>(in days) | Shipment timeliness<br>(in months) | Arrival<br>timeliness<br>(in months) | Delivery timeliness<br>(in months) |
| --- | --- | --- | --- | --- | --- |
| NTDeliver | 1.01 | 40.21 | 0.13 | 0.53 | 0.73 |
| p-value | (0.000) | (0.000) | (0.69) | (0.09) | (0.03) |
| Predicted direction | ↑ | ↓ | ↑ | ↑ | ↑ |
| Actual direction | ↑ | ↑ | ↑ NS | ↑ | ↑ |
| WHO region | Yes | Yes | Yes | Yes | Yes |
| Medicine type | Yes | Yes | Yes | Yes | Yes |
| Disease | Yes | Yes | Yes | Yes | Yes |
| Order size | Yes | Yes | Yes | Yes | Yes |
| Mode of shipment | Yes | No* | Yes | Yes | Yes |
| Observations | 1,052 | 249 | 636 | 783 | 636 |

NS = non-significant
\* Sample size too limited to incorporate variable

Notably, delivery timeliness, the KPI considered most important to measure supply chain performance, shows a positive and highly significant impact with a *p*-value of 0.03. The coefficient, 0.73, indicates a positive improvement of a little over half a month against the one-month pre-MDA benchmark. This result suggests information sharing is positively associated with earlier delivery of medicines to CMS. PO timeliness and arrival timeliness also demonstrate positive and significant results supporting the hypothesized direction, although arrival timeliness is significant at a lower confidence level. Information sharing appears to be strongly associated with a one-month improvement in PO timeliness, at the 99% confidence level. Arrival timeliness improvement by about half a month seems to also be associated with the information sharing, but with a *p*-value of 0.09 in the 90% confidence level, indicating a lower significance than the other performance indicators.

Conversely, the results for go signal timeliness appear to support the null hypothesis. While go signal timeliness is highly significant (p<0.01), even though the coefficient value is positive, this actually indicates a negative relationship with information sharing due to the calculation method for the variable (calculated as the difference between requested and actual go signal date). The coefficient suggests an increase in the difference between the go signal request and approval of about 40 days. Because there is no perceivable standard for establishing this request date and it is defined per request of the pharmaceutical manufacturer and any supporting partners, additional analysis provides further insight into how the go signal timeliness calculation may have changed after NTDeliver was implemented. Particularly, the pharmaceutical manufacturer may have started to provide earlier request dates, due to increased pressure to ensure on-time delivery after NTDeliver implementation.

These results provided indicate that the difference between PO date and go signal request date has significantly decreased, since the coefficient is negative (Table 3). This may indicate that since the implementation of information sharing through NTDeliver, go signal requests dates provided by the pharmaceutical manufacturer were earlier dates relative to the PO date. Also, the results indicate a significant reduction in the number of days between the go signal approved date and the shipment date since NTDeliver was implemented, indicating potentially that pharmaceutical manufacturers shipped medicines more quickly after the go signal was provided. Next, we study the impact of making country-level data publicly accessible and particularly promoting this data access to country program managers. Specifically, three sessions were conducted in 2018 to support publicizing the release NTD supply chain data to a country program manager audience: 1) an informational session at the program manager meeting in Kigali, Rwanda; 2) a webinar in the Pan American Health Organization (PAHO) region, and 3) a webinar hosted by ENVISION attended by country program managers from various regions. The hypothesis is that extending information sharing may have a positive impact for the following processes, because they are actively displayed in the public country pages, providing direct access for program managers to follow-up on status: shipment timeliness, arrival timeliness, and delivery timeliness. Table 4 provides the analysis results to answer the second research question regarding the impact of making country-level data publicly accessible.

**Table 3.** **Bivariate association between regression results showing the relationship between implementation of information sharing through NTDeliver and key differences between go signal dates**

| Dependent variable | Difference between PO date and go signal request date (in days) | Difference between go signal approved date and shipment date (in days) |
| --- | --- | --- |
| NTDeliver | - 48.86 | - 50.89 |
| p-value | 0.000 | 0.000 |
| WHO region | Yes | Yes |
| Medicine type | Yes | Yes |
| Disease | Yes | Yes |
| Order size | Yes | Yes |
| Mode of shipment | No* | No* |
| Observations | 263 | 263 |
| * Sample size too limited to incorporate variable |  |  |

**Table 4.** **Regression results showing the relationship between implementation of publicly accessible country-level data through NTDeliver and NTD supply chain key performance indicators**

| Dependent Variable | PO timeliness<br>(in months) | Go signal timeliness<br>(in days) | Shipment timeliness<br>(in months) | Arrival timeliness<br>(in months) | Delivery timeliness<br>(in months) |
| --- | --- | --- | --- | --- | --- |
| NTDeliver public access | 0.33 | - 14.88 | 2.57 | 2.88 | 2.82 |
| p-value | 0.49 | 0.42 | 0.001 | 0.003 | 0.01 |
| Predicted direction | NS | NS | ↑ | ↑ | ↑ |
| Actual direction | ↑NS | ↓NS | ↑ | ↑ | ↑ |
| WHO region | Yes | Yes | Yes | Yes | Yes |
| Medicine type | Yes | No* | Yes | Yes | Yes |
| Disease | Yes | Yes | Yes | Yes | Yes |
| Order size | Yes | Yes | Yes | Yes | Yes |
| Mode of shipment | Yes | No* | Yes | Yes | Yes |
| Observations | 387 | 132 | 296 | 262 | 255 |
| NS=non-significant |  |  |  |  |  |
| *Sample size too limited to incorporate variable |  |  |  |  |  |

The results show three variables with significant, positive correlation: shipment timeliness, arrival timeliness, and delivery timeliness—consistent with the hypothesis that these variables would be impacted since they are actively displayed in these publicly accessible pages. In the main regression results, shipment timeliness did not show any significance from the information sharing. In these results, shipment timeliness is the most significant variable at a *p*-value of 0.001 and with a substantial coefficient of 2.57, suggesting that while information sharing alone did not have any significant impact on this process, implementing the publicly accessible country pages appears to be impactful enough to significantly improve shipment timeliness by 2.5 months. For both arrival timeliness and delivery timeliness, which were also significant in the main regression results, the coefficients are substantially larger than in the main results and the *p*-values are substantially lower than those in the main regression results. Arrival timeliness in the main results was just on the cusp between being significant and non-significant, with a *p*-value of 0.09 in the 90% confidence level. In these results, arrival timeliness is significant with a *p*-value of 0.003. Also, the coefficient increases from 0.53 to 2.88, a marked improvement as well. Similarly, the important delivery timeliness performance indicator has a *p*-value of 0.011 in these results, compared to the main results in which the *p*-value was 0.03. The coefficient is 2.82, compared to the main results coefficient 0.73—indicating an improvement by over two months from the main results. These results suggest that while information sharing positively impacts arrival and delivery timeliness, increasing visibility of the country performance furthers the positive impact to the supply chain, particularly to downstream processes.

We also conducted further analyses to check the robustness of the results. First, we re-ran the regression accounting for a lag in impact from information sharing. The analysis revealed that that all dependent variables, except arrival timeliness, remain significant and generally consistent with the main results when accounting for a six-month theoretical “lag time to benefit,” considered as the time between implementing the intervention and observing improved outcomes.^27^ Additionally, we conducted a “double pretest” to test the validity of the “pre” group as comparison. This “double pretest” was used as a validity check to ensure that merely “history” and/or “maturation” is not the reason for the differences between the pretest and posttest results, rather than independent variable [28]. The pretest group was divided and compared for significant differences in performance to assure any differences in performance within the pretest group are minimal and/or less than the difference between the pretest and posttest groups. The groups were divided with roughly the same number of shipments in each group, with one group comprised of shipments with POs raised between 2006-2013 and the other with POs raised between 2014-2016 August. The results from this analysis indicated that only PO timeliness had a positive, significant difference between these two pretest groups. No other dependent variables have an observable difference in performance.

## Discussion

Lack of coordination and limited transparency are two top issues in humanitarian supply chains [29]. Although existing literature on for-profit supply chains suggests addressing the issues though information sharing, it is unclear whether and how information sharing can improve performance in the non-profit humanitarian context [10–13]. In this paper, we examined the impact of information sharing on humanitarian supply chain performance. This paper contributes to the small body of literature exploring humanitarian supply chain performance. Our analysis is the first to undertake an empirical study evaluating performance and information sharing both in the context of the NTD supply chain and in the broader humanitarian space. While most existing literature focuses on information sharing in for-profit supply chains, which is typically focused on sharing information between the buyer and supplier, this paper addresses the gap by investigating the impact of sharing information publicly for non-profit supply chains. The results of our study demonstrate the value of investment in supply chain performance measurement and information sharing towards the success of global health partnerships and such initiatives may be implemented in the broader context of humanitarian programs.

We find that information sharing is positively associated with the timeliness of several key stages in NTD supply chain, i.e., PO timeliness, arrival timeliness, and shipment timeliness, and consequently, the key success measure for the NTD supply chain—delivery timeliness. Furthermore, when detailed country-level data was shared with and promoted to NTD program managers in different countries, these timeliness measures improved even more. The impact of information sharing appears to be more significant when the information is released publicly and particularly promoted to country program managers, compared to when it is shared only with the supply chain partners (i.e., WHO and pharmaceutical donors). This observation may imply that it is more effective to share information with those accountable for successful outcomes and performance from the work, rather than only with those driving and directing the work.

The robustness checks also showed that even if the information sharing effect took time to make an impact, the performance still improved and can be attributed more confidently to implementation of NTDeliver. Furthermore, although there is no “control group” in our research design, we conduct a “double pretest” to investigate if other observed time-varying variables may contribute to the significant results. Comparison with performance before the information sharing was implemented suggests that the supply chain performance does not simply improve over time, with exception to one performance indicator. Only PO timeliness indicates a significant positive change over time during the period before NTDeliver was implemented. The reason for this change in PO timeliness may be attributed to a process change coinciding around the timing of the second pretest group defined in this analysis, which included medicines ordered from 2014 onward: the JAP was implemented by the WHO in 2013 to standardize processes to support an integrated application submission and review for donated medicines [23]. This JAP process most positively impacts the PO process as it promoted more coordination between the various levels of WHO offices to assure timely applications and order fulfillment for donated medicines [23].

This research has some limitations that may naturally inform future research. The quasi-experimental design used lacks a control group and random assignment since NTDeliver was implemented for all PC medicine donations managed through the NTD Supply Chain Forum [30]. Although we used a “double pretest” robustness check to verify our results, future research could study the impact of information sharing in a controlled setting. Furthermore, there is a growing desire for financial donors to understand the impact of investments from an outcomes perspective, especially in the interest of funding effective health innovations that offer value for money [31].While our research results certainly helps to validate the positive and measurable impact of the information sharing intervention on the supply chain, further research on how the supply chain performance improvements result in a reduction in delayed and/or missed MDAs would provide more perspectives on linking the delivery timeliness improvements to number of additional individuals reached. Lastly, with respect to the current global health climate, the COVID-19 pandemic had a significant impact on NTD programs, with the WHO recommending postponing MDAs to respect public health measures that advocate for physical distancing to slow the spread of the virus.^32^ Further research is needed to gain insight on new challenges from these disruptions to understand the impact by region and how information sharing may help to mitigate such disruption. Also, the data may potentially identify best practices that can provide insight on managing uncertainty for global health campaign supply chain planning during a pandemic.

Our results have important practical implications. As the global health aid landscape is becoming more focused on driving measurable performance and impact from investments, these findings support investing in supply chain systems and commitment to data transparency. Given the relationship between first mile supply chain performance and timeliness of MDAs, investment in supply chain information sharing is worthwhile to support improvements to NTD program management [7]. As the deadline approaches for achieving the 2030 targets set out in the new WHO roadmap and the relevant NTD goals in target 3.3 of the Sustainable Development Goals, there is a high degree of confidence that these results affirm that investment in supply chain information sharing is a critical to ensuring success. In fact, the new WHO roadmap dedicates an entire section to “Access and logistics,” in which supply chain management priorities for improvement are outlined under the umbrella concept that “effective supply chain management is vital to ensuring access to quality-assured NTD medicines and other products” [33]. Furthermore, the findings imply the benefits of information sharing are potentially maximized when extending information sharing to a broader audience, particularly focusing on program managers. Incorporating visibility to upstream data, such as attaching country applications or tracking the regional office approval date, may improve these processes as well. Such upstream processes have been noted as potentially impacting delivery timeliness by the WHO HQ—such as the fairly significant time taken for the regions to review applications—and further incorporation of these processes in NTDeliver may benefit the end-to-end NTD supply chain [7].

Given the significant volume of medicines and the number of people requiring these medicines, the research implications have the potential to impact global health programs affecting the health of tens to hundreds of millions of people. The research supports that, even in absence of financial remuneration, information sharing contributes measurable supply chain improvements and supports investing further in performance measurement in humanitarian supply chains. Beyond the NTD space, data transparency is generally viewed as a challenge with country governments citing national sovereignty and privacy in refusing to release data for public consumption [34]. Positive results from extending the information sharing argue in favor of the value and benefits of information sharing in the global health space. As the profile and importance of the supply chain continues to elevate in humanitarian programs, especially those in the healthcare space, there will be an opportunity to invest further in such performance measurement tools to bring more evidence-based approaches to decision making. This has significant potential to promote accountability and coordination resulting in goals achieved and improved health outcomes.

## Acknowledgments

The authors would like to acknowledge the NTD Supply Chain Forum for generously providing access to the data used in this research via NTDeliver. In particular, a big thank you is extended to key leaders of this Forum, including Tijana Williams, Dr. Christian Schröter, and Cassandra Holloway for support on granting this data access and disseminating results to the NTD supply chain partners of the Forum. We also thank TJ Muehleman and Emily Tunggala from the NTD Supply Chain Forum technology partner, Standard Co, for support to confirm details on the data sources in NTDeliver and additional insights on the data storage and management.

## Appendix

**Table A1.**
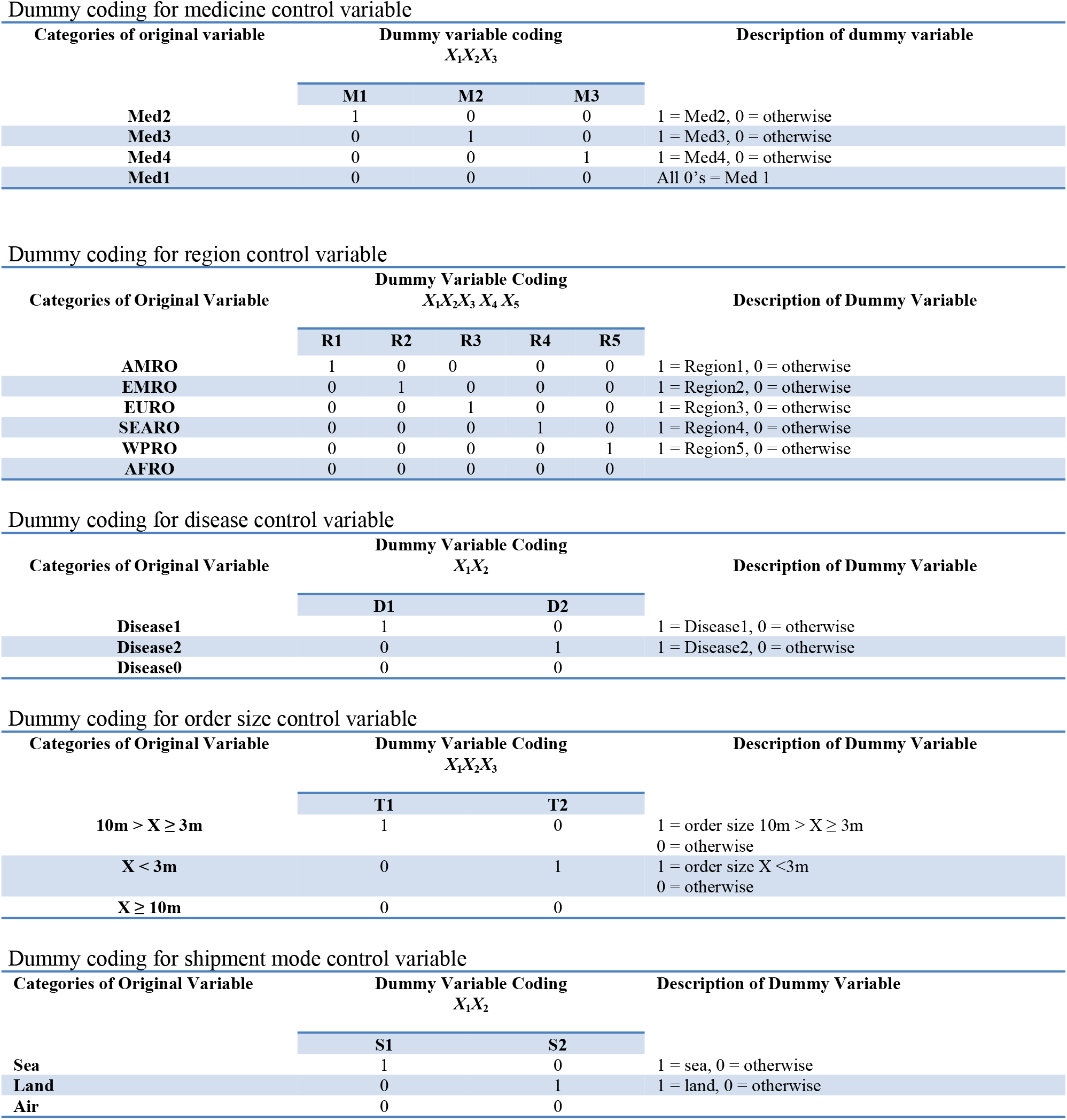
Dummy coding matrices.

**Table A2.**
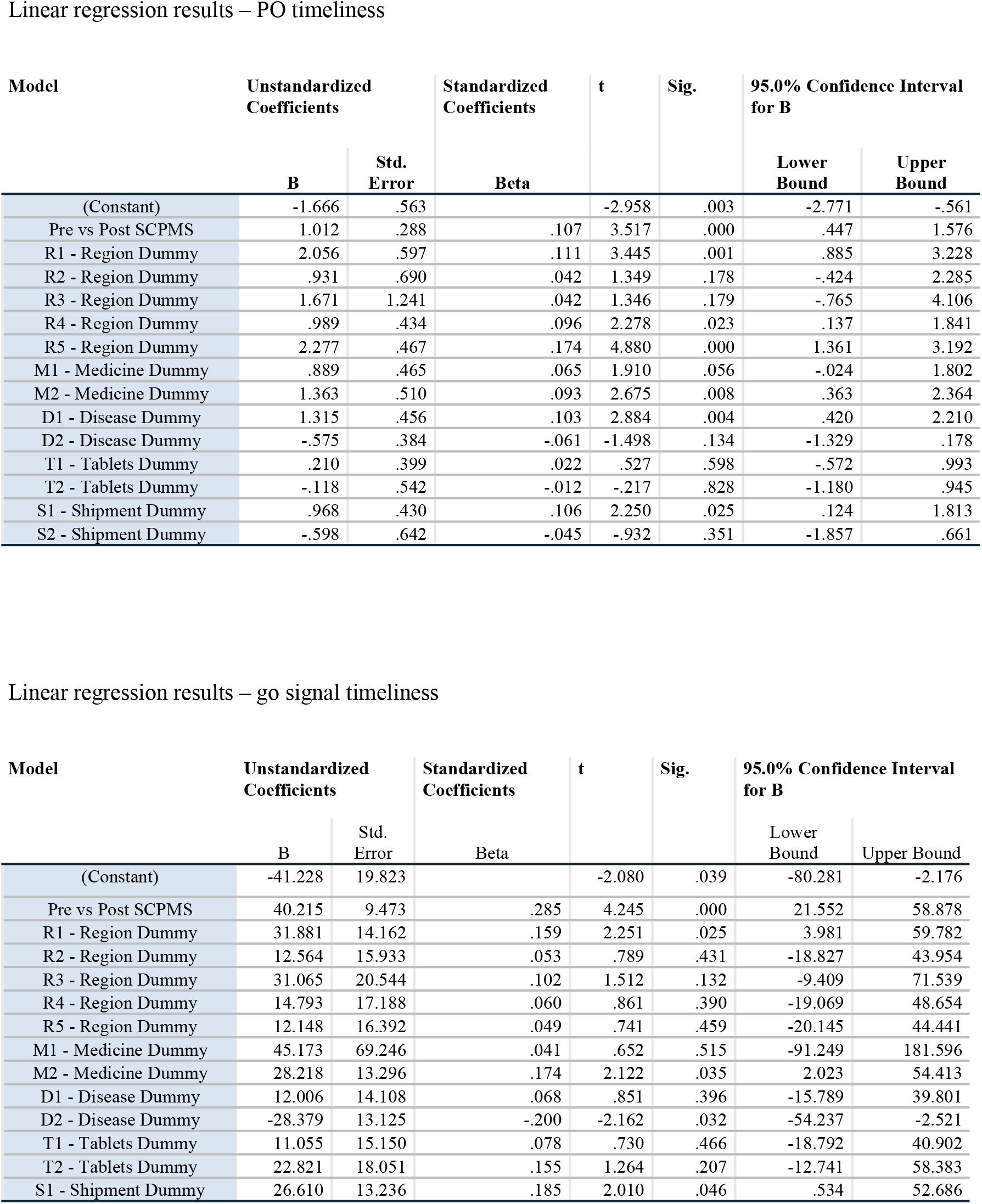
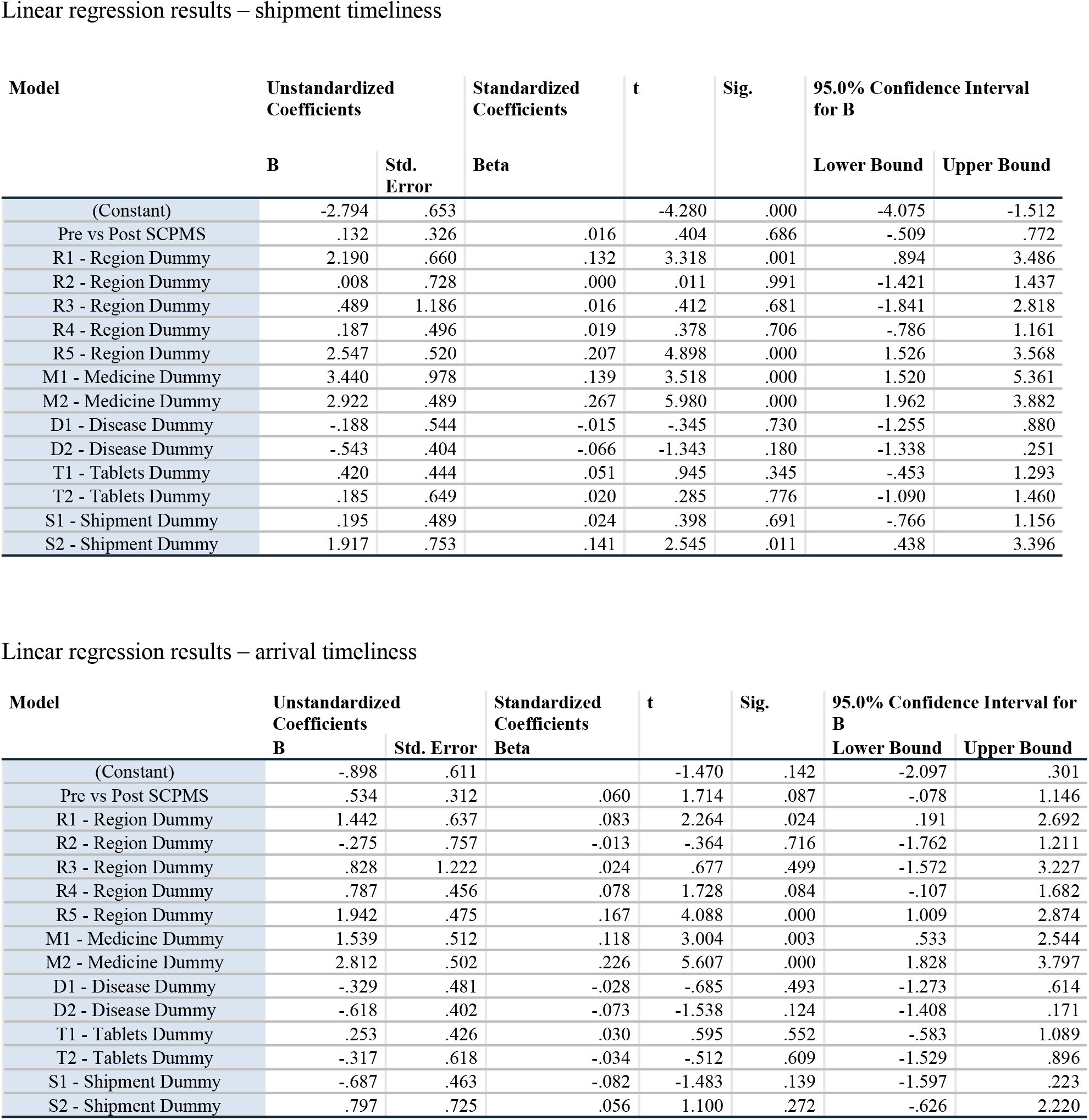
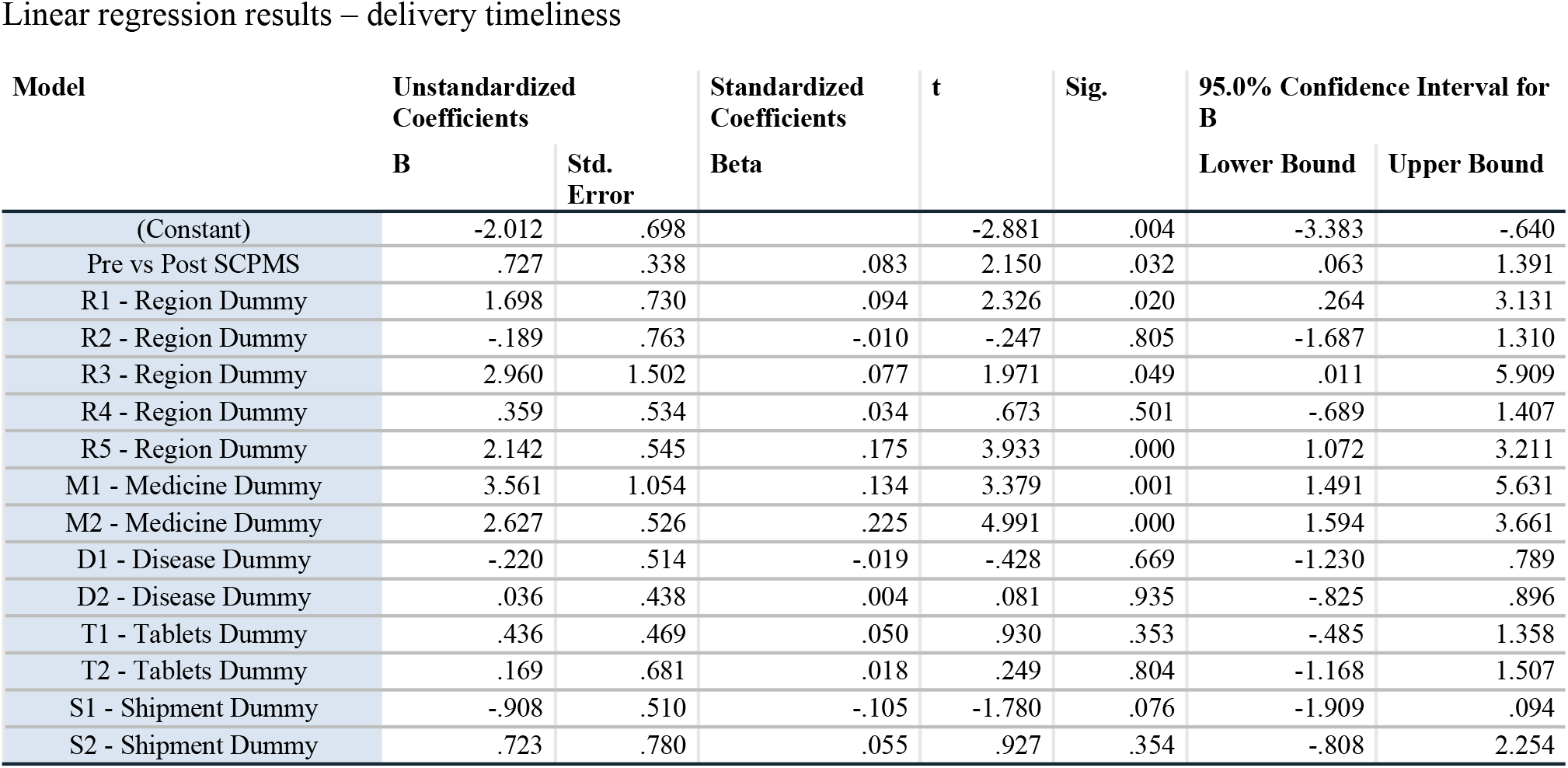
Full linear regression results.

**Table A3.** Robustness check results.

| Dependent Variable | PO timeliness (in months) | Go signal timeliness (in days) | Shipment timeliness (in months) | Arrival timeliness (in months) | Delivery timeliness (in months) |
| --- | --- | --- | --- | --- | --- |
| NTDeliver | 0.999 | 38.495 | - 0.026 | 0.204 | 0.741 |
| p-value | 0.002 | 0.000 | 0.937 | 0.553 | 0.046 |
| WHO region | Yes | Yes | Yes | Yes | Yes |
| Medicine type | Yes | Yes | Yes | Yes | Yes |
| Disease | Yes | Yes | Yes | Yes | Yes |
| Order size | Yes | Yes | Yes | Yes | Yes |
| Mode of shipment | Yes | No* | Yes | Yes | Yes |
| Observations | 970 | 226 | 569 | 715 | 567 |
| *Sample size too limited to incorporate variable |  |  |  |  |  |

Summary of ordinary least squares regression results applying a “double pretest”
| Dependent Variable | PO timeliness (in months) | Go signal timeliness (in days) | Shipment timeliness (in months) | Arrival timeliness (in months) | Delivery timeliness (in months) |
| --- | --- | --- | --- | --- | --- |
| NTDeliver | 1.42 | -30.476 | 0.991 | 0.581 | 0.173 |
| p-value | 0.000 | 0.293 | 0.215 | 0.203 | 0.818 |
| WHO region | Yes | Yes | Yes | Yes | Yes |
| Medicine type | Yes | Yes | Yes | Yes | Yes |
| Disease | Yes | Yes | Yes | Yes | Yes |
| Order size | Yes | Yes | Yes | Yes | Yes |
| Mode of shipment | Yes | No* | Yes | Yes | Yes |
| Observations | 664 | 117 | 339 | 520 | 380 |
| *Sample size too limited to incorporate variable |  |  |  |  |  |

